# The Impact of Excessive Muscle Co-contraction on Sit-to-Stand Performance in High-Heeled Footwear

**DOI:** 10.1101/2024.11.21.624701

**Authors:** Ganesh R. Naik, Amit N. Pujari

**Affiliations:** College of Medicine and Public Health, Flinders University, Adelaide, Australia; Design and Creative Technology Vertical, Torrens University, Wakefield Street, Adelaide SA 5000; Neu(RAL)^2^: NeuRAL Systems & Rehabilitation and Assistive Technologies Laboratory, School of Physics, Engineering and Computer Science, University of Hertfordshire, Hatfield, England, AL10 9AB; School of Engineering, University of Aberdeen, Aberdeen, Scotland, AB24 3UE

**Keywords:** Co-contraction, Co-contraction index, High heel shoes, Surface electromyography, Sit-to-stand, Vastus medialis, Vastus lateralis, Rectus femoris, Semitendinosus

## Abstract

This study aimed to analyse the effects of co-contraction on quadriceps and hamstring muscles during sit-to-stand tasks for females wearing shoes with different heel heights. The study aimed to identify compensatory strategies during sit to stand task in response to excessive muscle co-contraction during high heeled gait. Sixteen healthy young women (age: 24.4±1.7 years, body mass index: 18.4±1kg/m2, weight: 50.2±5.2kg, height: 1.63±4.4m) participated in this study. Electromyography signals were recorded from three quadriceps (vastus medialis, vastus lateralis, and rectus femoris) and one hamstring (semitendinosus) muscles. The participants wore shoes with different heights, including 4cm, 6cm, 8cm, and 10cm. For each heel height, we computed co-contraction index to measure postural balance using quadriceps to hamstring muscle pairs. The results that were obtained and quantified with statistical measures show that for elevated shoes if co-contraction increases, both quadriceps and hamstring muscles tend to compensate. This suggests that the capacity of quadriceps and hamstring muscles to compensate is essential to retain normal walking and sit-to-stand (STS) tasks in co-contracted persons. However, the compensation mechanisms may induce imbalance, muscle stiffness, and fatigue for women who regularly use high-heeled shoes during sit-to-stand tasks.

## 1. Introduction

Nowadays, women regularly wear high-heeled shoes (HHS) for day to day activities which include, walking, stair ascent, stair descent, sit-to-stand (STS), and stand-to-sit-returning (STSR) tasks. Regular use of HHS has been reported to negatively impact different body structures, change gait mechanics, and result in musculoskeletal problems (Cronin, 2014; Mika et al., 2013). STS and STSR tasks are the most frequently performed activities in daily life (Bolink et al., 2012; Dehail et al., 2007). These tasks are described as a motion of the human body from a stable sitting-down position to a straight-up-standing position and vice versa (Kim et al., 2011; Roebroeck et al., 1994). These tasks require higher muscle power and coordination in the balance system than other daily tasks, such as walking and stair climbing (Abe et al., 2010; van Der Kruk et al., 2021). This demands posture adjustments and optimal neuromuscular coordination of the quadriceps and hamstring muscles (Bolink et al., 2012; Buckley et al., 2009).

Co-contraction, the synchronized activation of agonist and antagonist muscles (antagonistic pairs), occurs in several daily events, including postural control, walking, and running (Kellis & Kouvelioti, 2009; Wang & Gutierrez-Farewik, 2014; Weir et al., 1998). Busse et al. (2005), report that co-contraction is the mechanism that regulates the simultaneous activity of agonist and antagonist muscles crossing the same joint. Other research shows that excessive co-contraction can cause inefficient or abnormal movements in some neuromuscular pathologies and is even associated with normal aging (Kellis et al., 2003; Palmieri-Smith et al., 2009). Excessive or poorly controlled co-contraction is reported to be a major cause of inefficient gait (Thomas et al., 2022) in individuals with cerebral palsy (Lundh et al., 2018), with potentially negative repercussions on the quality of life (Thomas et al., 2022) such as restricting joint motion and increasing energy expenditure, in individuals with CP (Gharehbolagh et al., 2023; Unnithan et al., 1996). As it has been posited that agonist-antagonist muscle co-contraction reflects a deliberate neural control strategy to preserve effector-level control allowing stabilizing motor actions without having to control individual muscles separately (Latash, 2018); investigating co-contraction during regular/daily tasks (Kellis & Kouvelioti, 2009; Wang & Gutierrez-Farewik, 2014; Weir et al., 1998), as well as during specific contractions can provide valuable insights about changing muscle behaviours and neural control strategy (Pujari et al., 2019).

Due to increased mechanical demands associated with STS and STSR tasks, it is reasonable to expect activation of the lower extremity muscles to increase with gait speed (Janssen et al., 2002; Lord et al., 2002; Roy et al., 2007). Prior research shows that women appear to preferentially activate the lateral quadriceps and hamstring muscles during STS while simultaneously displaying less medial thigh muscle activation (Palmieri-Smith et al., 2009). Moreover, as quoted by Lloyd et al., the quadriceps and hamstring muscles have the potential to provide dynamic knee stability because of their abduction and/or adduction moments (Heiden et al., 2009; Lloyd & Buchanan, 2001). To maintain joint stability and body balance during high loading tasks of STS and STSR, it is crucial that muscle co-contraction of the involved muscles is optimum, as co-contraction is known to be used to maintain joint stability, whereas excessive co-contraction is a result of pathological changes in neuromuscular changes. To our knowledge, effects of muscle co-contraction of HHS for STS have not been examined.

Regularly wearing HHS alters the neuromechanics of walking, compromises muscle efficiency, causes discomfort, and increases the risk of strain injuries (Cronin, 2014). Additionally, it has been postulated that HHS may contribute to developing and progressing knee osteoarthritis (OA) (Edwards et al., 2008; Kim et al., 2011). As wearing HHS can not only lead to abnormal alterations in neuromuscular and musculoskeletal behavior but also contribute to long-term complications such as osteoarthritis, this study attempts to analyze activation and co-contraction patterns of quadriceps and hamstring agonist/antagonist muscle pairs to investigate potential neuromuscular alterations caused by HHS. We specifically focus on the Rectus femoris (RF), Vastus lateralis (VL), Vastus medialis (VM) and Semitendinosus (ST) during STS of different HHS gait. Two hypotheses were tested in this study. First, we hypothesized that elevated heels would have greater co-contraction of the quadriceps to hamstring muscles during STS tasks with HHS. The second hypothesis was that elevated heel height will be associated with increased muscle activity in the quadriceps and hamstrings. This may lead to altered joint kinematics and kinetics during the sit-to-stand task.

While the ankle joint, stabilised by the anterior tibialis and gastrocnemius muscles, is significantly affected by high-heeled footwear, this study focused on the knee joint and the co-contraction of the quadriceps and hamstring muscles. This decision was based on several considerations. Firstly, the knee joint is subjected to increased stress and altered biomechanics due to the changes in weight distribution and joint angles associated with high-heeled shoes. Secondly, co-contraction of the quadriceps and hamstrings is a common strategy employed by the body to enhance joint stability and control movement (Lu & Chang, 2012; Winter, 2009). By analysing the co-contraction patterns in this muscle pair, we aimed to gain insights into the compensatory mechanisms adopted by individuals to maintain balance and stability during the sit-to-stand task while wearing high-heeled shoes.

## 2 Materials and Methods

An exploratory repeated measures study was conducted using data collected from young female participants for the proposed research. The materials and methods used for this study are explained below.

### 2.1 Participants

The University of Technology Sydney, Sydney (UTS), Human Research Ethics Committee approved the experimental protocol (Ethics details: UTS HREC 2013000728) for this study and adheres to the declaration of Helsinki. Sixteen healthy young women participated in this study (age: 19.4±1.7 years, body mass index: 21.4±1kg/m^2^, weight: 50.2±5.2kg, height: 1.63±0.1m). As a pilot study, this study was based on convenience sample (of sixteen women) and no power calculations were done. All the participants had no musculoskeletal disorders or injuries of the lower extremity, were not pregnant, and did not have a history of surgery on the lower extremity. An information sheet was given, and all participants signed a written consent form in presence of the researcher before the experiment. The participants were familiar with wearing HHS.

### 2.4 Study design

For this study, shoes with four different heel heights were chosen, including 4cm, 6cm, 8cm, and 10cm. The shoes used for the experiment are shown in Figure 1 (a). The surface of the heels is approximately 1cm^2^ for all shoes, defined as a stiletto in the fashion industry. To maintain control, the shape and style of these shoes are chosen to be as similar as possible.

**Figure 1.**
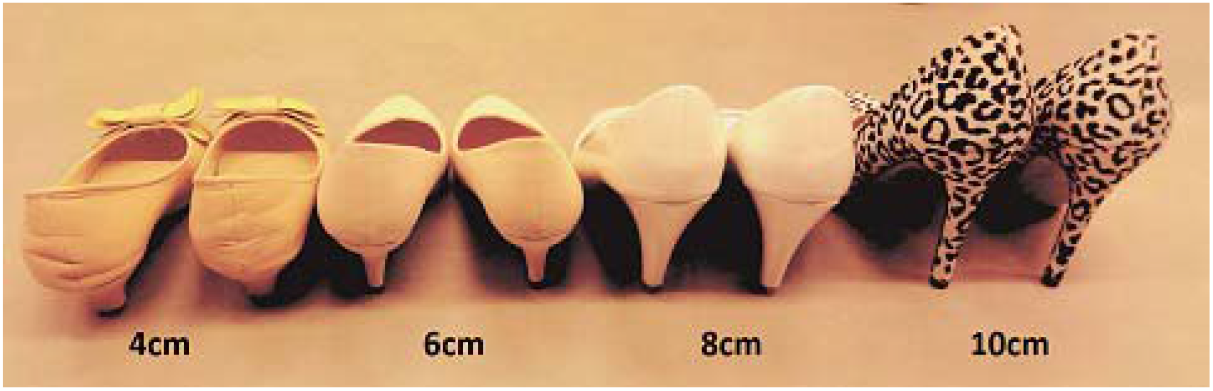
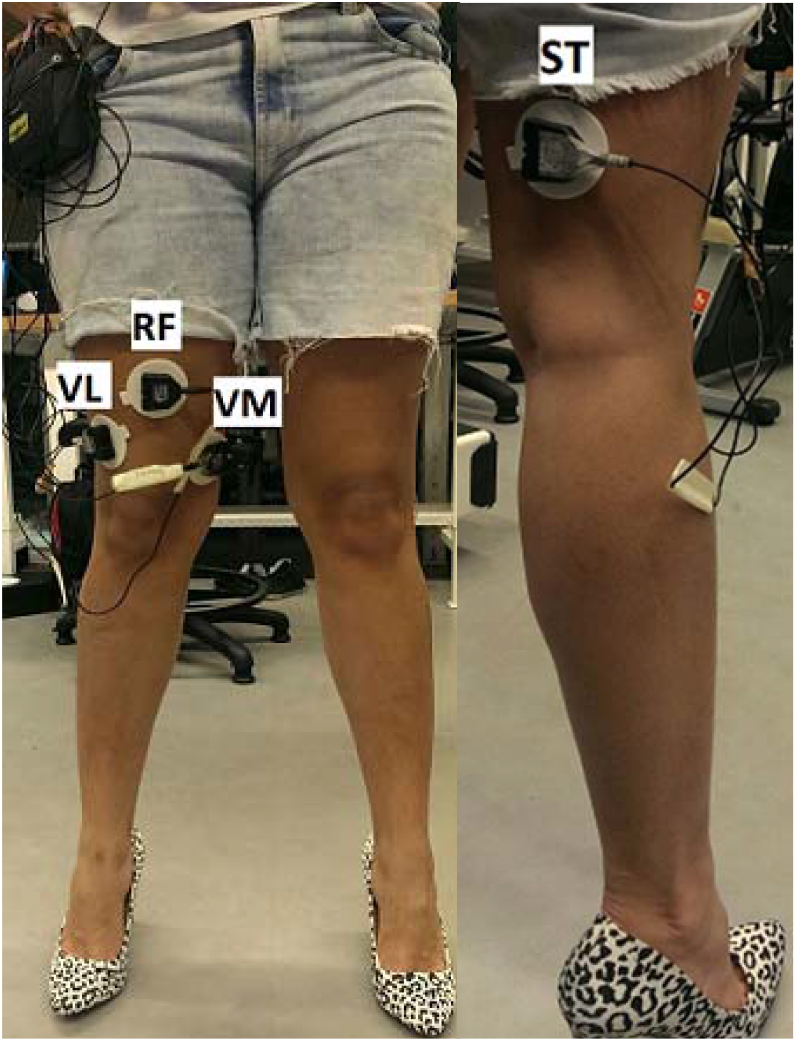
a) Shoes used for the experiment. b) Electrode locations for the quadriceps (left) and hamstring (right) muscles.

Participants were not habitual high-heel wearers. Prior to the experiment, they were introduced to high-heeled shoes of varying heights. To minimise pre-experimental muscle strain, participants tried on all shoe heights without weight-bearing. High-heeled shoes were worn exclusively during the experimental period

### 2.3 Data Acquisition

The sEMG signals were recorded from three quadriceps muscles and a hamstring muscle, which include RF, VM, VL and ST muscles. The quadriceps and hamstring muscle groups, specifically the VM and ST, were selected for this study. These muscles play crucial roles in knee extension and flexion, respectively, and have been extensively studied in relation to lower limb biomechanics and functional tasks such as sit-to-stand (Pan et al., 2023; Sadeh et al., 2023). While the biceps femoris also contributes to knee flexion, the semitendinosus was chosen due to its larger muscle mass and its potential for greater influence on joint moments and power generation during the task (Onishi et al., 2002). The electrodes were placed on the dominant leg. The electrode locations used for the experiment are shown in Figure 1 (b). To identify the dominant leg, participants were asked which leg they would choose to kick a ball with, and the chosen leg was considered the dominant one. The placement of electrodes was configured according to the SENIAM guidelines (Hermens et al., 1999). The electrodes were silver-silver triode with a fixed inter-electrode distance of 2 cm and a diameter of 1cm (Thought Technology, Montreal, Quebec, Canada). Prior to electrode placement, hair was removed from the skin surface, and the skin was cleaned with an alcohol wipe to reduce skin impedance. The skin was then allowed to air dry before electrode placement to ensure optimal signal quality. The sEMG signal was recorded by Flexcomp Infiniti encoder system and transmitted wirelessly to a computer through Bluetooth (Biograph Infiniti) at 2048 samples/sec.

All tasks were performed in a health technology laboratory. The sequence of wearing different heights of shoes was randomly assigned. Detailed explanations of the planned experiments were given before data recording so that participants could familiarize themselves with the environment and procedure. For the STS task, participants sat in an armless chair. Participants were instructed to sit on a standard chair with a seat height of 46 cm with their feet flat on the floor. They were asked to put their arm across their chest. Thus, the arms would not be used to assist the movement of standing up. Their feet were placed in a set position so that movement of feet and legs was not required when they stood up from their seated position. After the participants settled in the chair, they sat still for five seconds and were signaled to stand up by the word “stand.” Participants were required to remain standing for five seconds until they heard the word “sit.” Participants carried out three repetitions of a sit to stand task under each of four conditions (of wearing 4 cm, 6 cm, 8 cm, and 10 cm high heels). The sit and stand period duration was maintained across all conditions (experiments). After each condition (i.e. After wearing 4cm HHS, then 6cm HHS, then 8cm HHL, etc.), about 3 to 5 minutes rest was given to avoid potential muscle fatigue.

### 2.4 Data Processing

Data were exported from Biograph Infiniti and then processed using MATLAB R2017a software. sEMG data was normalised using maximal voluntary isometric contraction (MVIC) value (recorded from the same muscle) as the reference value (Halaki & Ginn, 2012). For that, a sEMG signal from a given muscle was used to the sEMG recorded from the same muscle during MVIC as the reference value. A 4^th^ order Butterworth band pass filter with a frequency range of 20 to 450 Hz was applied to reject any frequency outside this range, likely to be noise (Potvin & Brown, 2004; Zhang & Zhou, 2013). Due to movement artifacts in the initial and final transient phases of the test, the signals generated during these periods (i.e., before 5% and after 95% of the total time of the test) were discarded.

#### 2.4.1 Quantification of co-contraction

Muscle co-contraction was assessed between the quadriceps (RF, VL, and VM) and hamstring (ST) muscle pairs, which include VL – ST, VM – ST and RF – ST combinations. The co-contraction index (CCI) is computed by integrating the ratio of the linear-enveloped EMG multiplied by the sum of the EMG magnitudes for the quadriceps and hamstring muscle pairs as described by Nelson-Wong et al. (2012).

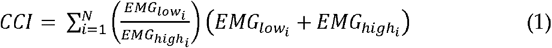

where *N* is the total number of data points for the time frame of interest, *EMG*_*lowi*_ is the lower EMG value at the ith data point, and *EMG*_*highi*_ is the higher EMG value at the ith data point. The CCI provided a measure of relative activation of the muscle pairs at each instance of the gait cycle (Hubley-Kozey et al., 2008; Lewek et al., 2004). Also, larger and smaller CCI values represent greater and lower muscle co-contractions respectively (Tsai et al., 2012). During the analysis, Equation 1 was applied for the sEMG data of each muscle pair (VL-ST, VM-ST, and RF-ST) that was recorded for a 5-s duration. Since sEMG data were sampled at 2,048 Hz, there were 10,240 data points included in each 5-s window (*N*). For each *i*th point within the 5-s window, the linear-enveloped EMG magnitudes were compared by taking the low over the high value ratio and then multiplied by the sum of the two magnitudes (Equation 1). These products were then summed over the 10,240 data points comprising the 5-s window. The resulting values for 10240 data points are averaged to produce a single value representing the co-contraction index for each muscle pair during one STS task. The overall results for 3 STS trials were then averaged for statistical analysis. The above procedure was repeated for all HHS tasks and the 16 subjects.

To assess the normality of the data, the Shapiro-Wilk test was used. As the data was found to be normally distributed, parametric statistical tests (i.e., t-tests, ANOVA) were employed to analyse the data. A repeated measure analysis of variance (ANOVA) test (using MATLAB software) was performed to compare the CCI calculated for each of the three muscle pairs (RF-ST, VM-ST, and VL-ST) using four different heel heights. The statistical level of significance was fixed at p<0.05 (95% confidence intervals). The effect size (small, medium, or large) of all co-contraction parameters was assessed using Cohen’s d (standardized mean differences). Taking into account the cut-off established by Cohen, the effect size can be small (∼0.2), medium (∼0.5), or large (∼0.8) (Cohen, 1992).

## 3. Results

Pooled (mean and standard deviation) CCI from the four heel heights for the three muscle group combinations is shown in Figure 2. The highest CCI ratio was found for the VM-ST muscle pairs, while the lowest values were found for RF-ST and VL-ST variants. The results indicated that the CCI ratios increased for elevated HHS; this is because both quadriceps and hamstring muscles exert higher abduction and/or adduction moments for high-level muscle activities (Lloyd & Buchanan, 2001; Palmieri-Smith et al., 2009). Also, due to increased mechanical demands associated with STS tasks, it is reasonable to expect activation of the lower extremity muscles, especially quadriceps and hamstring muscles, to increase with HHS (Seyedali et al., 2012). Predominantly, the simultaneous recruitment of muscles that produce moments in opposite directions, as happens during increased antagonistic muscle co-contraction around the joint, has a substantial influence on the movement patterns of the knee during STS tasks (Sirin & Patla, 1987).

**Figure 2.**
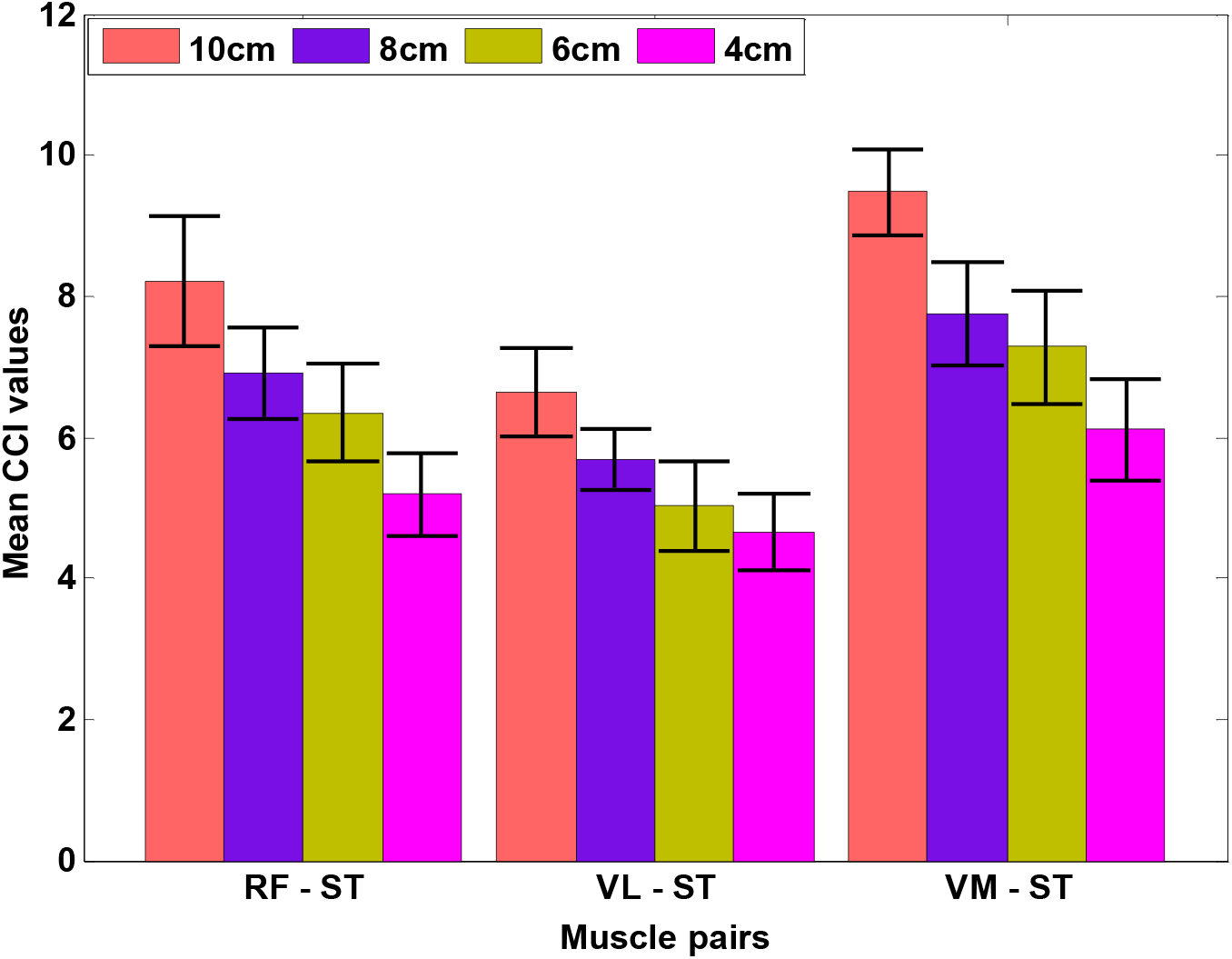
The mean and standard deviation of Co-contraction Index (CCI) from the four heel heights for three muscle group combinations. RF: Rectus Femoris, VL: Vastus Lateralis, VM: Vastus Medialis, ST: Semitendinosus

The distributions of muscle involvement (quadriceps and hamstring) in the different HHS-STS tasks, as measured by the CCI ratio (percentage of each muscle pair), are shown in Figure 3. From the results, it is interesting to see that RF-ST, VL-ST, and VM-ST showed similar CCI distribution (approximately 34%, 28%, and 38%, respectively) irrespective of heel height. In other words, the CCI distribution remains constant for RF-ST, VL-ST, and VM-ST for all HHS. This is very likely because both the quadriceps and hamstring muscles have the potential to provide dynamic frontal-plane knee stability and have the capacity to balance variable abduction-adduction loads (Lloyd & Buchanan, 2001; Palmieri-Smith et al., 2009).

**Figure 3.**
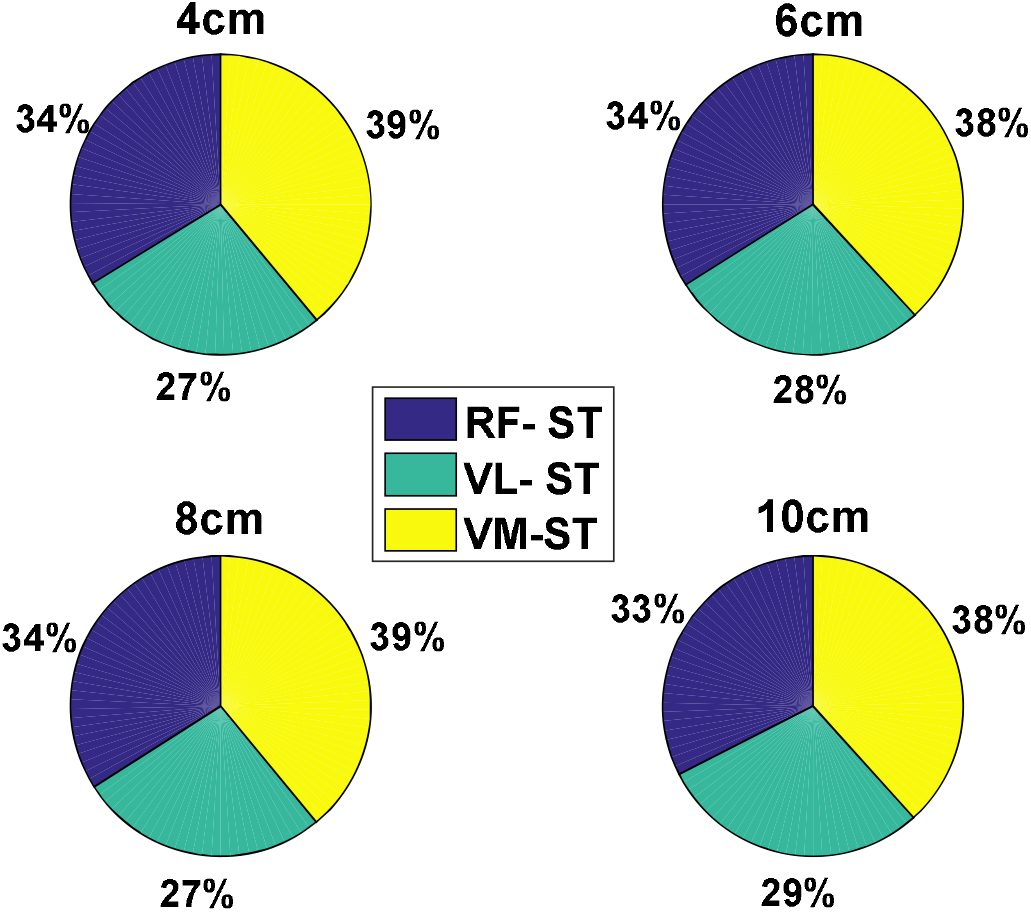
The CCI distribution (%) for RF-ST, VL-ST, and VM-ST for all heel heights. RF: Rectus Femoris, VL: Vastus Lateralis, VM: Vastus Medialis, ST: Semitendinosus

For the RF-ST co-contraction, no significant differences were found between 8cm and 6cm HHS (*p*>0.05; Cohen’s d = 0.84), as shown in Table 1. The RF-ST co-contraction index was highest for 10cm HHS and lowest for 4cm HHS. There were significant differences in CCI values between 10cm-8cm, 10cm-6cm, 10cm-4cm, 8cm-4cm, and 6cm-4cm groups for RF-ST co-contraction during STS tasks (*p*<0.05). For the VL-ST co-contraction, no significant differences were found between 6cm and 4cm (*p*>0.05; Cohen’s d = 0.64) HHS (Table 2). The VL-ST co-contraction index was highest for 10cm HHS and lowest for 4cm HHS. Significant differences in CCI values were found between 10cm-8cm, 10cm-6cm, 10cm-4cm, 8cm-6cm, and 8cm-4cm groups for VL-ST co-contraction during STS tasks (*p*<0.05). For the VM-ST co-contraction (similar to RF-ST), no significant differences were found between 8cm and 6cm HHS (*p*>0.05; Cohen’s d = 0.62), as shown in Table 3. The VM-ST co-contraction index was highest for 10cm HHS and lowest for 4cm HHS. There were significant differences in CCI values between 10cm-8cm, 10cm-6cm, 10cm-4cm, 8cm-4cm and 6cm-4cm groups for VL-ST co-contraction during STS tasks (*p*<0.05). The sEMG signal patterns for the VL-ST, RF-ST, and VM-ST for one of the subjects are shown in Figures 4 – 6, respectively. Comparable sEMG patterns (Figures 4-6) were observed for all the participants. From the plots, it is evident that a similar trend of sEMG patterns can be seen for the RF-ST, 6-8cm; VL-ST, 4-6cm and VM-ST, 6-8cm results. The results show that VL-ST pair (refer to Figure 4) exhibits similar properties for 4cm-6cm heel heights, whereas RF-ST and VM-ST pairs (refer to Figures 5-6) show similar results for 6cm-8cm heel heights. The results are also justified with high Cohen’s index values indicating the similarity among the mentioned HHS pairs. Similar findings are seen in Remaud et al. (2009), which show similar muscle activities for VM and RF muscles, whereas specific muscle activity was observed for VL muscle for isotonic and isokinetic contractions.

**Table 1:**
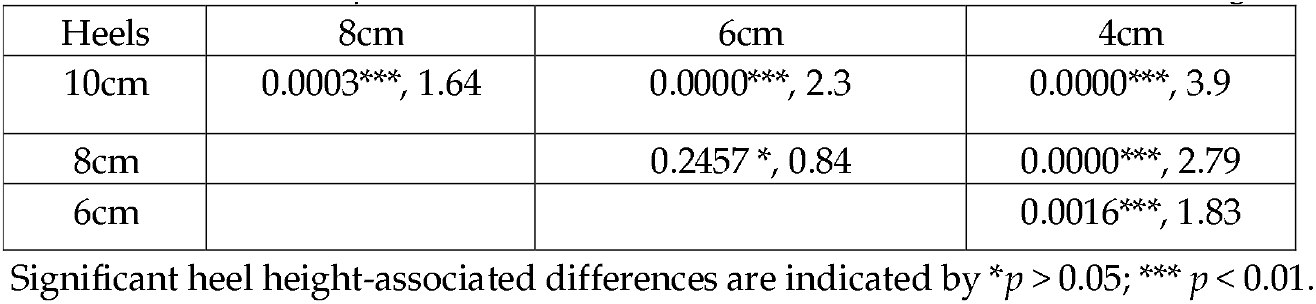
Pairwise ANOVA (*p* - values) and Cohen’s d effect size for all four heel heights using RF-ST.

| Heels | 8cm | 6cm | 4cm |
| --- | --- | --- | --- |
| 10cm | 0.0003***, 1.64 | 0.0000***, 2.3 | 0.0000***, 3.9 |
| 8cm |  | 0.2457*, 0.84 | 0.0000***, 2.79 |
| 6cm |  |  | 0.0016***, 1.83 |

Significant heel height-associated differences are indicated by \* $p > 0.05$ ; \*\*\* $p < 0.01$ .

**Table 2:** Pairwise ANOVA (*p* - values) and Cohen’s d effect size for all four heel heights using VL-ST Heels 8cm 6cm 4cm.

| Heels | 8cm | 6cm | 4cm |
| --- | --- | --- | --- |
| 10cm | 0.0009***, 1.77 | 0.0000***, 2.53 | 0.0000***, 3.41 |
| 8cm |  | 0.0316*, 1.21 | 0.0003***, 2.17 |
| 6cm |  |  | 0.3824*, 0.64 |

Significant heel height-associated differences are indicated by \* $p > 0.05$ ; \*\*\* $p < 0.01$ .

**Table 3:** Pairwise ANOVA (*p* - values) and Cohen’s d effect size for all four heel heights using VM-ST.

| Heels | 8cm | 6cm | 4cm |
| --- | --- | --- | --- |
| 10cm | 0.0000***, 2.53 | 0.0000***, 3.11 | 0.0000***, 5.03 |
| 8cm |  | 0.3591*, 0.62 | 0.0000***, 2.24 |
| 6cm |  |  | 0.0014***, 1.53 |

Significant heel height-associated differences are indicated by \* $p > 0.05$ ; \*\*\* $p < 0.01$ .

**Figure 4.**
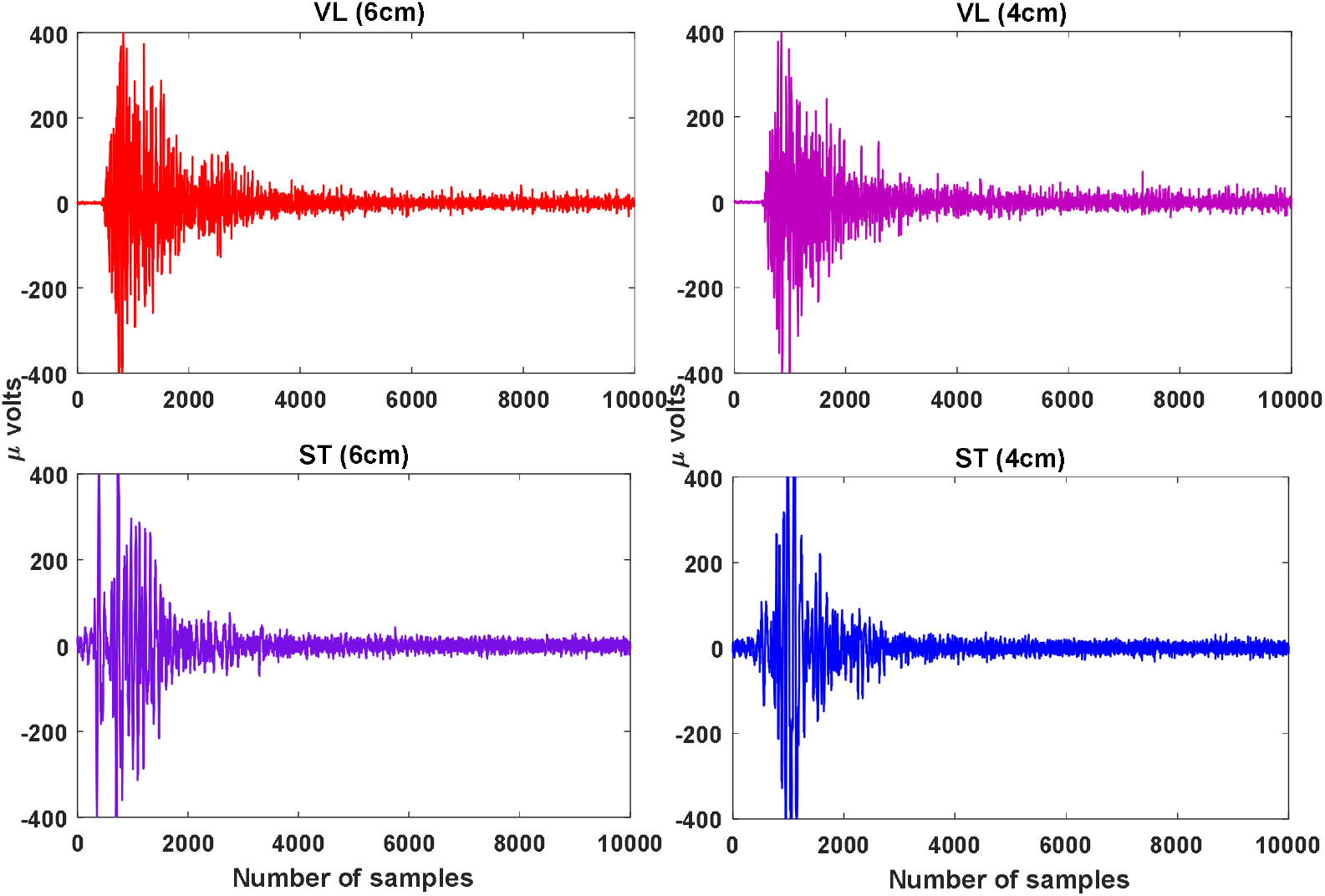
The raw sEMG patterns for VL and ST muscles for 6cm and 8cm. Here, the x-axis represents the samples and y-axis represent the amplitude in micro (μ) volts. VL: Vastus Lateralis, ST: Semitendinosus

**Figure 5.**
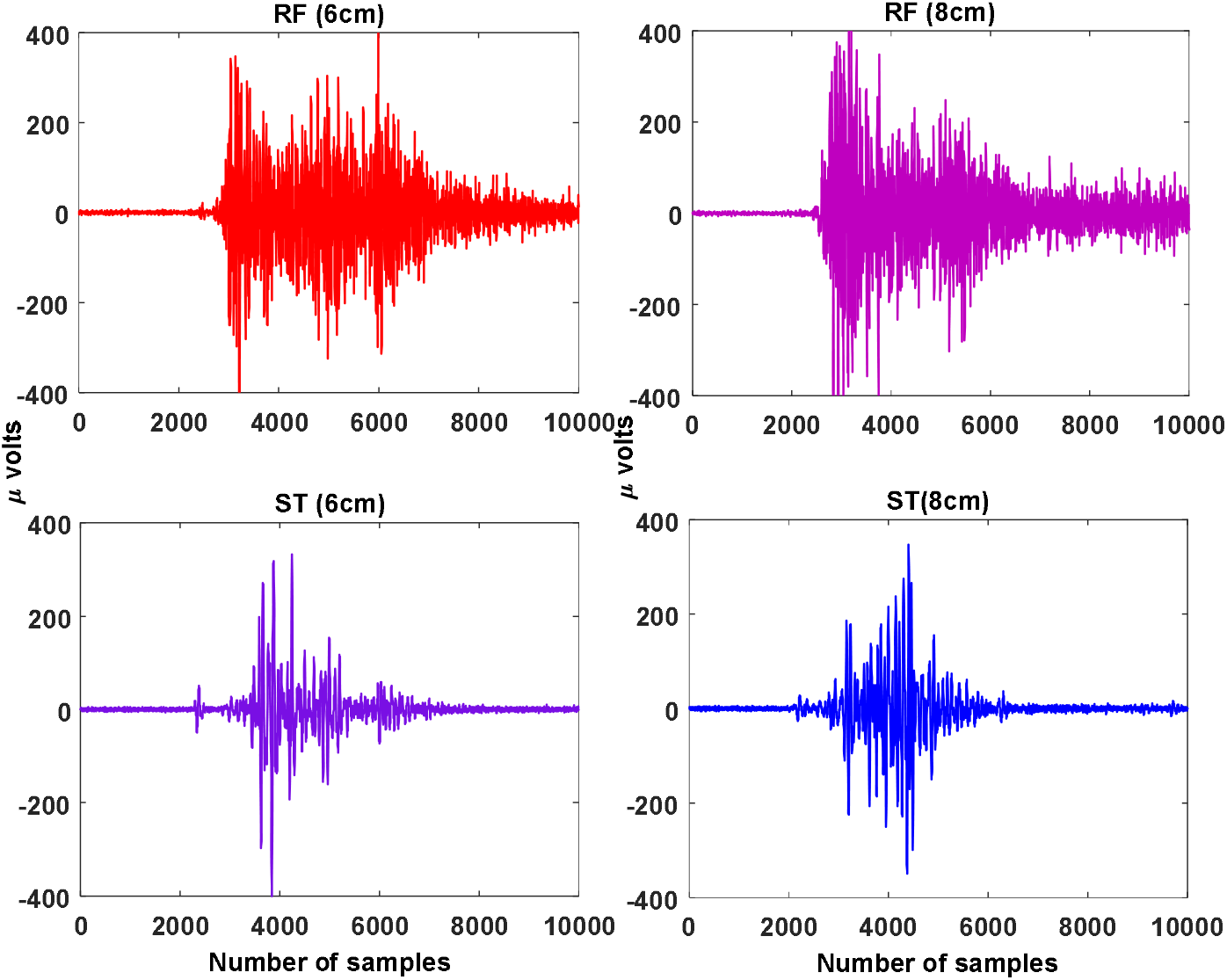
The raw sEMG patterns for RF and ST muscles for 6cm and 8cm. Here, the x-axis represents the samples and y-axis represent the amplitude in micro (μ) volts. RF: Rectus Femoris, ST: Semitendinosus

**Figure 6.**
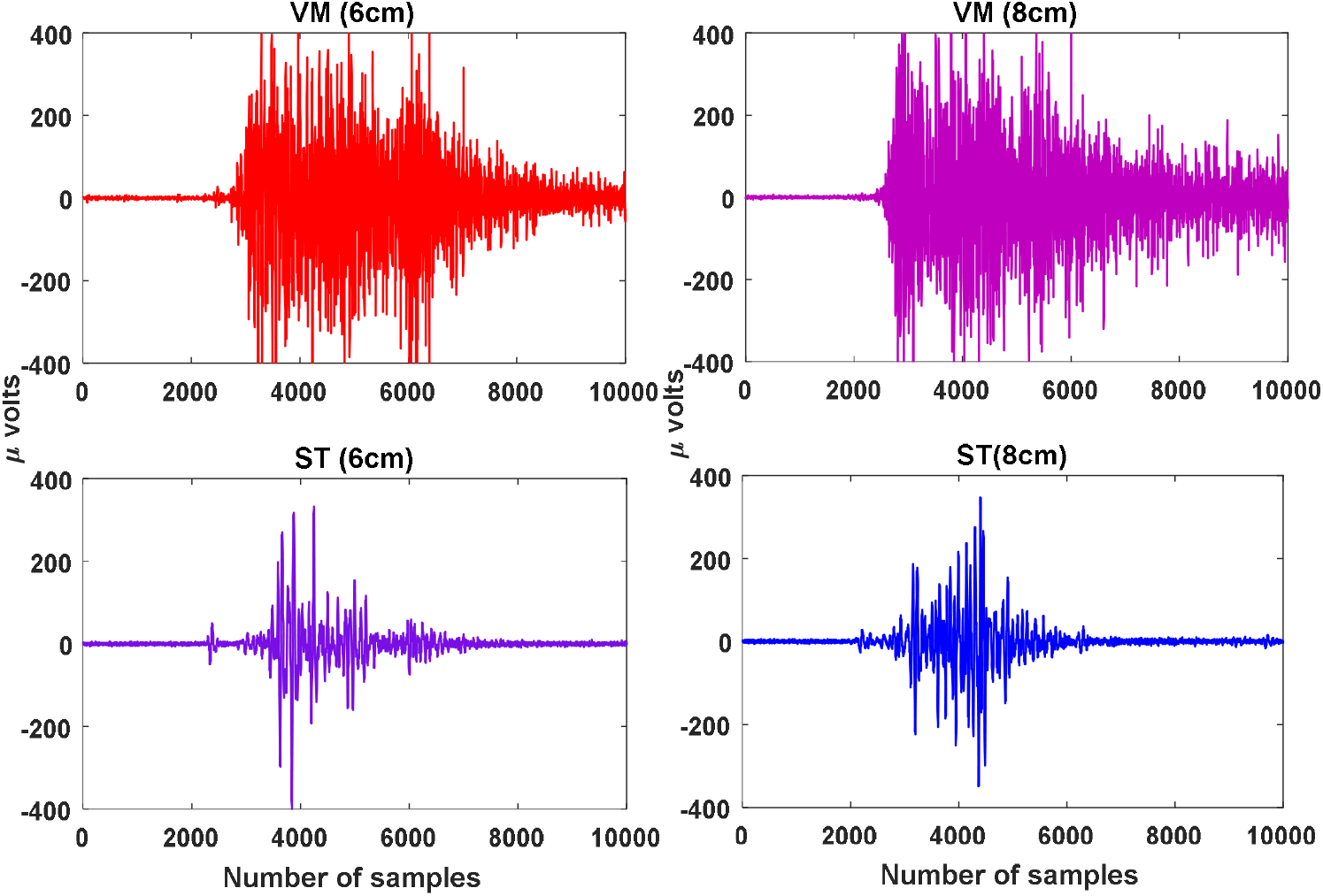
The raw sEMG patterns for VM and ST muscles for 6cm and 8cm. Here, the x-axis represents the samples and y-axis represent the amplitude in micro (μ) volts. VM: Vastus Medialis, ST: Semitendinosus

## 4. Discussion

The major findings of this study are that: 1) irrespective of subjects, in 6cm-8cm heel height, compensation (or adjustment) needed to maintain balance or stability between Antagonistic muscle pairs RF and ST occur similarly with different co-contraction values. 2) For 6cm-8cm heel height, compensation (or adjustment) to maintain balance or stability between Antagonistic muscle pairs VM and ST occurs similarly with different co-contraction values. 3) For 4cm-6cm heel height, compensation (or adjustment) to maintain balance or stability between Antagonistic muscle pairs VL and ST occurs similarly with different co-contraction values.

Understanding how quadriceps and hamstring muscles behave during STS can help clarify the motor control strategies exerted (during STS tasks) using muscle activation patterns to overcome excessive muscle co-contraction. Goulart and Valls-Solé (2001) reported the differential action of related muscles during the STS task and determined that the pattern of muscle activity remained constant when the initially seated posture changed (participant seated in different places). As stated by Seyedali et al. (2012) co-contractions may represent a limb stiffening strategy to enhance stability during phases of initial heel strike for HHS, which may result in increasing CCI values for elevated HHS.

The CCI of quadriceps to hamstring ratio remained the same for all four heel heights (Figure 3). Hypothetically, the net effect of the contribution provided by both quadriceps and hamstring muscles should be approximately constant under different co-contraction levels, since CCI ratio remains the same for all HHS heights. Hence, it could be expected that elevated HHS exerts more external work to maintain the same quadriceps to hamstring ratio compared to lower HHS. Mainly, elevated HHS movements theoretically require the muscle to perform more work on a given STS compared to lower HHS movements. Moreover, if co-contraction increases, both quadriceps and hamstring muscles can compensate for elevated shoes. The above results are also in agreement with the previous study by Wang and Gutierrez-Farewik (2014), which states that due to muscle redundancy, various neuro-motor strategies may exist to compensate for excessive muscle co-contraction.

STS tasks demand complex and optimum neuromuscular coordination and postural changes to control the body and prevent loss of balance (Bishop et al., 2005; Janssen et al., 2002). According to Dehail *et al*. (Dehail *et al*., 2007), the human body must make necessary adjustments to maintain postural balance. One such scenario, related to the STS task where significant modifications or essential adjustments are needed is wearing high-heeled shoes. Barton et al. (2009), reported that regular usage of high heel shoes for STS and related tasks might contribute to changes in body posture, and may induce low back pain in women.

Our research results indicated that capacity of the quadriceps and hamstring muscles to compensate is fundamental for retaining normal STS tasks with higher muscle co-contraction for elevated HHS. However, we can argue that women appear to co-contract their muscles to enhance stability and support during STS tasks with HHS. Women may employ co-contraction strategies during STS tasks to stabilize and provide extra shock absorption during heel strike.

The increased muscle co-contraction observed in this study during the sit-to-stand task in high-heeled footwear suggests that individuals may employ compensatory strategies to maintain balance and stability. High heels significantly alter the body’s center of mass and the distribution of weight on the feet, leading to increased stress on joints, particularly the knees and ankles. To mitigate these effects, individuals may increase co-contraction of agonist and antagonist muscles around the knee and ankle joints, thereby enhancing joint stability and shock absorption, especially during heel strike. Furthermore, the reduced ankle dorsiflexion imposed by high heels may necessitate increased muscle activity to maintain balance and propulsion (Cronin, 2014; Hapsari & Xiong, 2016).

### 4.1 Theoretical Contribution

This study contributes to the growing body of research on the biomechanical effects of high-heeled footwear. By demonstrating increased muscle co-contraction during the sit-to-stand task, this research provides further evidence for the compensatory strategies employed by individuals to maintain balance and stability in challenging footwear conditions. These findings offer valuable insights into the neuromuscular mechanisms underlying gait adaptation and the potential impact of high-heeled footwear on musculoskeletal health.

### 4.2 Practical Contribution

The findings of this study have practical implications for healthcare professionals, footwear designers, and individuals who frequently wear high-heeled shoes. Healthcare providers can use this information to educate patients about the potential risks associated with high-heel wear, such as increased muscle strain and joint stress. Footwear designers can leverage these findings to develop footwear that minimizes the negative biomechanical effects of high heels, such as by incorporating innovative designs that promote better foot alignment and shock absorption. Individuals who frequently wear high heels can benefit from understanding the potential consequences and consider limiting their use or choosing footwear with lower heels to reduce the risk of musculoskeletal injuries.

### 4.3 Limitations

While this study provides valuable insights into the biomechanical effects of high-heeled footwear, it is important to acknowledge certain limitations. Firstly, the sample size was relatively small, which may have limited the statistical power of the study. Future research with larger sample sizes can provide more robust evidence. Secondly, the study focused on a specific population of healthy young women, and the results may not be generalizable to other populations, such as older adults or individuals with specific musculoskeletal conditions. Future studies should consider a more diverse range of participants to assess the impact of high-heeled footwear on different populations.

Future research could also explore the long-term effects of high-heel wear on musculoskeletal health, including the development of chronic pain conditions such as plantar fasciitis and osteoarthritis. Additionally, investigating the impact of different heel heights and shoe styles on muscle activation patterns and joint loading would provide further insights into the mechanisms underlying the adverse effects of high-heeled footwear. By addressing these limitations and exploring these research directions, we can gain a more comprehensive understanding of the biomechanical consequences of high-heel wear and develop strategies to mitigate the associated risks.

## 5. Conclusion

The findings of this study have revealed significant differences in quadriceps – hamstring muscle co-contraction for HHS during STS tasks. Additionally, the lower and higher heel shoes had significant differences in co-contraction levels of the quadriceps and hamstring musculature. The occurrence of co-contractions depends on the phase of movements, along with the demands and characteristics of the muscle during STS tasks.

This exploratory study aimed to quantify the effect of co-contraction for HHS. The results support the hypothesis that quadriceps to hamstring co-contraction increases for elevated HHS. Our study findings indicated that capacity of the quadriceps and hamstring muscles to compensate is fundamental for retaining normal STS tasks with muscle co-contraction. From the results, it could be expected that elevated HHS exerts more external work to maintain the same quadriceps-to-hamstring ratio compared to lower HHS. Hence, the compensation mechanisms used by lower limb muscles may induce imbalance, muscle stiffness, and fatigue with regular usage of high heel shoes in women during STS task.

There are some inherent limitations in this study. The STS task conditions may have been too similar to reveal differences in co-contraction. Future efforts should examine the effect of these factors on residual limb activation and co-contraction patterns.

## Author Contributions

GRN conceptualized the study and completed experiments/data collection. GRN and ANP completed data signal processing and analysis (validation, visualisation). GRN prepared original draft and ANP carried out reviewing, editing. Both GRN and ANP prepared and agreed upon the final version of the manuscript.

## Data availability

All data underlying the findings will be are available from authors upon request.

